# Metabolites with benzene ring from sugarcane leaf play important role in plant-*Spodoptera frugiperda* interaction

**DOI:** 10.1101/2023.08.29.555346

**Authors:** Liangyinan Su, Chunyu Hu, Chaoqi Wang, Baoshan Chen, Yang Zhao

**Author notes:** Corresponding author: Yang Zhao; Baoshan Chen.

## Abstract

Secondary metabolism plays important role in plant growth and development, however, the relationship between secondary metabolism and adaptive plant-insect communication is largely unknown. The present study used sugarcane line highly susceptible to *Spodoptera frugiperda* and sister line with medium resistance to analyze the role of plant non-volatile organic compounds (NOCs) and volatile organic compounds (VOCs) in sugarcane-*S. frugiperda* interaction. A total of 46 plant NOCs and 15 plant VOCs significantly different between resistant and susceptible lines and were continuously up-regulated and down-regulated at different time points before/after *S*. *frugiperda* treatment were screened. Phenolic acids containing benzene ring accounted for the largest proportion of differential NOCs. Levels of 66.7% of these phenolic acids were higher in susceptible line. Feeding supplemented with NOCs showed that phenoxyacetic acid (phenolic acid) and 4-methoxybenzaldehyde (aromatic phenolic acid) both increased the male-to-female ratio of *S. frugiperda*. Aromatics containing benzene ring, accounted for the largest of differential VOCs in susceptible line. Two aromatics, *p*-cymene and benzene and 1-ethenyl-4-methoxy-, with higher level in susceptible line, were attractive to *S. frugiperda*. Terpenoids, aldehyde, and esters accounted for most of higher-in-resistant VOCs, with most tested to be repellent to *S. frugiperda*. Furthermore, transcriptome analysis of *S. frugiperda* feeding on susceptible and resistant lines combined with feeding assays revealed that tryptophan, as a precursor of aromatic compounds that also contains benzene ring, could promote the growth and development of *S. frugiperda* in nutritional deficiency condition. These findings together suggested that benzene-ring containing compounds play a critical role in plant-*Spodoptera frugiperda* interaction.

## Introduction

Plant secondary metabolites constitute an important tool for researching plant-herbivore interactions (Bilas et al., 2021). According to the properties (i.e., gases, liquids, and solids) of secondary metabolites, they can be divided into volatile organic compounds (VOCs) and non-volatile organic compounds (NOCs). The NOCs are important compounds for host plants to defend against herbivore insects, and the NOCs related to plant defense against herbivore insects are mainly phenolic acids and flavonoids. The accumulation and emission of plant VOCs in plant tissues may alter plant-insect interactions and act as attractants or deterrents for insects (Szucs et al., 2011). The role of secondary metabolism in adaptive plant-insect communication is still largely unknown.

Plant NOCs can affect the growth and development of herbivore insects. Ruuhola et al. found that the higher concentration of phenolic acids in *Salix* spp. can significantly reduce the growth rate of *Operophtera brumata* larvae (Ruuhola et al., 2001). Flavonoids in cotton, such as quercetin and rutin, can inhibit the growth and development of *Pectinophora gossypiella* and *Heliothis virescens* (Shaver and Lukefahr, 1969). Another two flavonoids, isoorientin and isoorientin-7-O-arabinosyl-glucoside, could inhibit the growth, development and reproduction of *Sitobion avenae* (Liu et al., 2003).

Plant VOCs can affect insect host selection (Divekar et al., 2022). By analyzing the effects of tobacco HIPVs in insect behavioral preference tests, De Moraes et al. suggested that female moths (*Heliothis virescens*) can recognize HIPVs and disregard already infected plants, thus avoiding competition with other Lepidoptera moths and/or otherwise upregulating defenses of tobacco (De Moraes et al., 2001). Brouce et al. found that the HIPV (*E*, *E*) –4, 8, 12-trimethyltrideca-1, 3, 7, 11-tetraene (TMTT) from *Arabidopsis thaliana* had repellent effects on *Myzus persicae* (Bruce et al., 2008).

Most recent studies on plant-insect interactions through secondary metabolites have focused on model crops (De Moraes et al., 2001; Bodenhausen and Reymond, 2007; Riedlmeier et al., 2017; Yuan et al., 2017), few studies have been conducted on other graminaceous crops. Lepidoptera insects are one of the main pests of graminaceous crops. Damage caused by lepidoptera insect infestation during the seedling stage can affect sugarcane photosynthesis, making stalks vulnerable to lodging and, in severe cases, death, leading to reduced yield (Souza et al., 2021). The fall armyworm *Spodoptera frugiperda* (J. E. Smith, 1797) (Lepidoptera: Noctuidae) is one of the most important agricultural pests worldwide (Montezano et al., 2018), which mainly harms corn, rice, sugarcane and other crops. *S. frugiperda* can selectively bite heart lobes and new leaves of the host plant in the early stages, causing serious impacts on the major crop industries (Chormule et al., 2019). To date, interaction between sugarcane (*Saccharum officinarum*) and *S. frugiperda* has not been fully demonstrated. Based on field investigations, we found that sugarcane lines of hybrid offspring had different susceptibilities to *S. frugiperda*, therefore hypothesized the secondary metabolites in resistant and susceptible sugarcane lines may play an important role in sugarcane-*S. frugiperda* interaction. In order to investigate the differences in secondary metabolites, we performed analyses before and after *S. frugiperda* infestations using resistant and susceptible sugarcane lines, and verified the effects of differential secondary metabolites as well as an amino acid precursor tryptophan on the growth, development and behavior of *S. frugiperda*. These findings shed light on the possible interaction mechanisms of graminaceous crops and *S. frugiperda*.

## Results

### Differences in plant NOCs between resistant and susceptible sugarcane lines

We found a highly *S. frugiperda* – susceptible sugarcane line (S-115) and a medium *S. frugiperda* – resistant line (R-111), which were obtained from the same genetic population (**Figure S1** and **Table S1**). We speculated that the differential secondary metabolites in resistant and susceptible lines might play an important role in the interaction between sugarcane and *S. frugiperda*. In order to explore the differences of NOCs in resistant and susceptible sugarcane lines, NOCs were measured using UPLC-MS/MS at 0h, 1h, 4h, and 8h time points before/after *S. frugiperda* infestation (**Table S2** and **Figure S2-S3**). We selected 453 NOCs with matching rating 3 (the compound is consistent with the database compound) for subsequent analyses, excluding compounds which may have been introduced during the sampling or metabolite detection processes.

The largest proportion of 453 NOCs was phenolic acids (**Table S3**), accounting for 17.8 % and 17.6 % respectively in both healthy resistant and susceptible lines. Notably however, levels of NOCs were greatly lower in susceptible line. A total of 168 differential NOCs were identified in uninfected healthy plants, with the relative amount of 57 NOCs higher in resistant line and 111 NOCs higher in susceptible line (**Table S4**).

Oral secretions of *S. frugiperda* combined with mechanical damage that mimicked *S. frugiperda* infection were treated on sugarcane leaves, which triggered differential plant NOC changes. In the resistant line, levels of lipids increased rapidly, followed by phenolic acids, alkaloids and other compounds. At 1 h and 4h treatment, phenolic acids constituted the largest proportion of differential NOCs in the resistant line, about 20.7 % and 22.4 %, respectively. At 8 h treatment, the proportion of amino acids derivatives and alkaloids similarly increased to largest proportion with 12.5 % of differential NOCs (**Table S5**). In susceptible line, levels of lipid and phenolic acids rapidly increased following treatment. At 1 h treatment, similarly, phenolic acids constituted the largest proportion of differential NOCs for 17.6 %. At 4 h and 8 h, the amounts of flavonoids increased to the largest, for 21.6 % and 28.1 % of differential NOCs, respectively (**Table S6**).

In order to select NOCs that may related to the response to *S. frugiperda* infection, the continuously up-regulated or down-regulated differential NOCs were selected (**Figures 1**). The 34 differential NOCs (27 upregulated and 7 downregulated) were selected in resistant line (**Figures 1A, B**), while 17 differential NOCs (8 upregulated and 9 downregulated) were selected in susceptible line (**Figures 1C, D**). Excluding repeated differential NOCs, we finally selected 46 compounds, belonging to 11 categories (**Table 1** and **Figures 1E**). The relative amounts of most phenolic acids from 46 compounds were up-regulated after treatment in both resistant and susceptible lines, except phenoxyacetic acid that was detected only in healthy resistant line and 4-methoxybenzaldehyde detected only in healthy susceptible line. Similarly, we also found that apigenin (belonging to flavonoids) and 12-hydroxyoctadecanoic acid (belonging to lipids) were detected only in healthy susceptible line (**Figures 1F** and **Table S7**).

**Figure 1.**
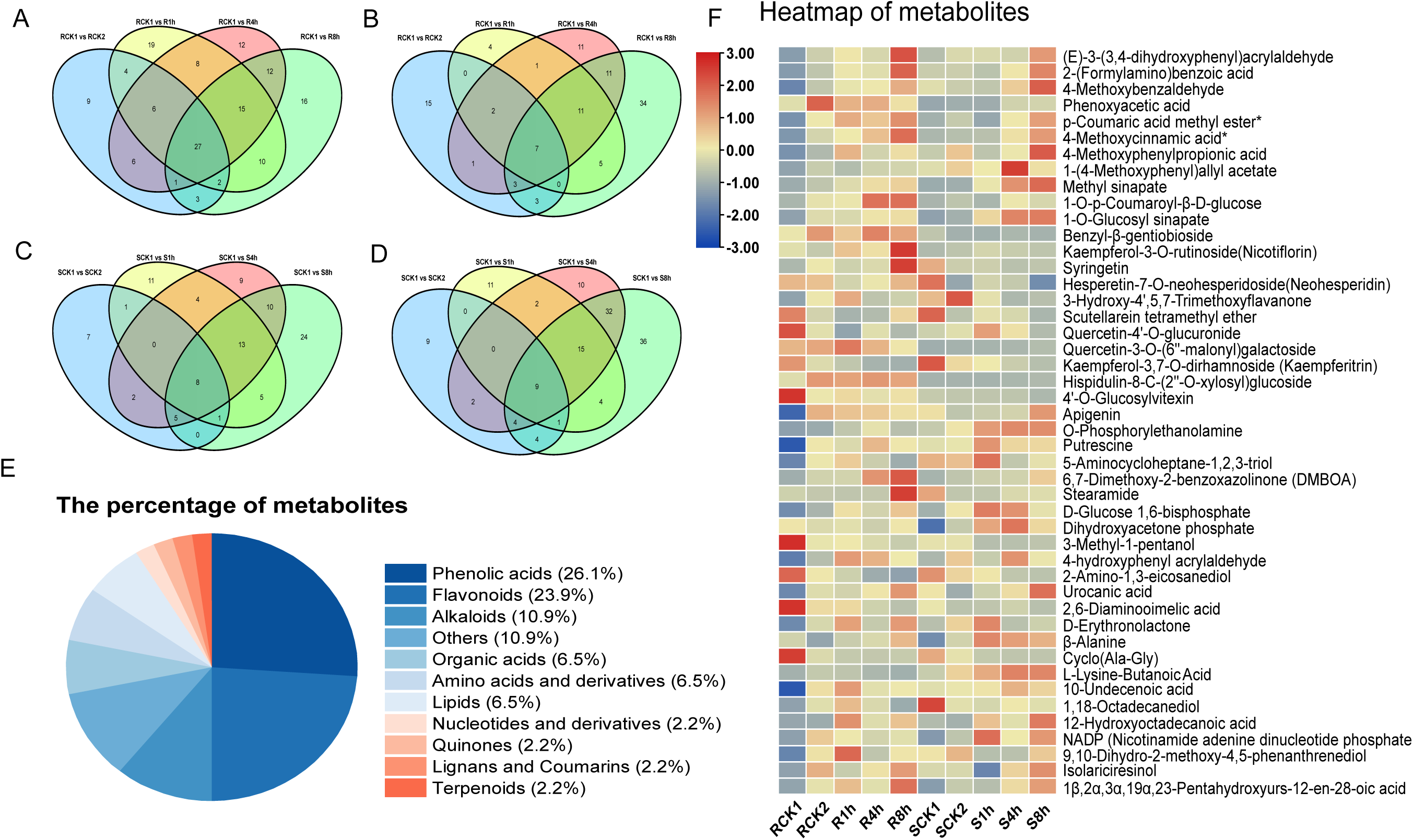
Analysis of NOCs in resistant and susceptible sugarcane lines. (A) The number of NOCs that were continuously up-regulated at different time points in resistant line (B) The number of NOCs that were continuously down-regulated at different time points in resistant line. (C) The number of NOCs that were continuously up-regulated at different time points in susceptible line. (D) The number of NOCs that were continuously down-regulated at different time points in susceptible line. Resistant group: untreated (RCK1), mechanical damage for one hour (RCK2); mechanical damage and oral secretion treatment for one hour (R1h); mechanical damage and oral secretion treatment for four hours (R4h); mechanical damage and oral secretion treatment for eight hours (R8h). Susceptible group: untreated (SCK1); mechanical damage for one hour (SCK2); mechanical damage and oral secretion treatment for one hour (S1h); mechanical damage and oral secretion treatment for four hours (S4h); mechanical damage and oral secretion treatment for eight hours (S8h). (E) Proportion of compounds in selected 46 NOCs. (F) Heatmap of 46 NOCs levels in different treatment of resistant and susceptible sugarcane lines.

**Table 1.** Information of selected compounds based on plant NOC analysis.

| Compound information |  |  | Relative volatilization amount (105)* |  |  |  |  |
| --- | --- | --- | --- | --- | --- | --- | --- |
| Class I | Compounds |  | CK1 | CK2 | 1h† | 4h‡ | 8h§ |
| Phenolic acids | (E)-3-(3,4-dihydroxyphenyl)acrylaldehyde | R-111 | 1.52 | 3.24 | 4.28 | 3.49 | 8.17 |
|  |  | S-115 | 2.67 | 3.59 | 3.58 | 3.34 | 6.27 |
| Phenolic acids | 2-(Formylamino)benzoic acid | R-111 | 4.71 | 10.52 | 15.05 | 13.83 | 30.08 |
|  |  | S-115 | 11.51 | 11.71 | 10.10 | 16.02 | 26.14 |
| Phenolic acids | 4-Methoxybenzaldehyde | R-111 | 0.00 | 0.53 | 0.76 | 0.80 | 1.19 |
|  |  | S-115 | 0.71 | 0.58 | 0.53 | 1.00 | 1.69 |
| Phenolic acids | Phenoxyacetic acid | R-111 | 0.43 | 1.37 | 0.87 | 0.90 | 0.58 |
|  |  | S-115 | 0.00 | 0.05 | 0.04 | 0.34 | 0.35 |
| Phenolic acids | *p*-Coumaric acid methyl ester* | R-111 | 0.20 | 0.56 | 0.77 | 0.68 | 0.86 |
|  |  | S-115 | 0.33 | 0.44 | 0.29 | 0.57 | 0.81 |
| Phenolic acids | 4-Methoxycinnamic acid* | R-111 | 0.44 | 1.12 | 1.34 | 1.69 | 2.39 |
|  |  | S-115 | 0.80 | 0.88 | 0.91 | 1.41 | 2.01 |
| Phenolic acids | 4-Methoxyphenylpropionic acid | R-111 | 0.15 | 0.47 | 1.19 | 0.67 | 0.94 |
|  |  | S-115 | 0.61 | 1.08 | 0.55 | 0.84 | 1.75 |
| Phenolic acids | 1-(4-Methoxyphenyl)allyl acetate | R-111 | 0.06 | 0.21 | 0.16 | 0.23 | 0.24 |
|  |  | S-115 | 0.27 | 0.40 | 0.32 | 0.82 | 0.35 |
| Phenolic acids | Methyl sinapate | R-111 | 0.27 | 0.66 | 0.99 | 1.36 | 1.50 |
|  |  | S-115 | 0.26 | 0.31 | 1.13 | 2.23 | 2.72 |
| Phenolic acids | 1-O-*p*-Coumaroyl-β-D-glucose | R-111 | 0.72 | 1.63 | 1.70 | 3.68 | 3.73 |
|  |  | S-115 | 0.71 | 1.02 | 0.72 | 1.74 | 1.39 |
| Phenolic acids | 1-O-Glucosyl sinapate | R-111 | 11.88 | 45.82 | 44.06 | 57.05 | 60.03 |
|  |  | S-115 | 12.41 | 20.54 | 63.02 | 99.61 | 103.40 |
| Phenolic acids | Benzyl-β-gentiobioside | R-111 | 0.19 | 0.41 | 0.29 | 0.46 | 0.37 |
|  |  | S-115 | 0.09 | 0.03 | 0.02 | 0.04 | 0.03 |
| Flavonoids | Kaempferol-3-O-rutinoside(Nicotiflorin) | R-111 | 0.12 | 0.05 | 0.25 | 0.18 | 0.60 |
|  |  | S-115 | 0.00 | 0.03 | 0.09 | 0.06 | 0.03 |
| Flavonoids | Syringetin | R-111 | 0.03 | 0.15 | 0.09 | 0.12 | 0.50 |
|  |  | S-115 | 0.27 | 0.09 | 0.09 | 0.05 | 0.06 |
| Flavonoids | Hesperetin-7-O-neohesperidoside(Neohesperidin) | R-111 | 0.54 | 0.51 | 0.32 | 0.38 | 0.54 |
|  |  | S-115 | 0.73 | 0.16 | 0.32 | 0.28 | 0.06 |
| Flavonoids | 3-Hydroxy-4',5,7-Trimethoxyflavone | R-111 | 0.03 | 0.09 | 0.14 | 0.06 | 0.07 |
|  |  | S-115 | 0.12 | 0.19 | 0.08 | 0.05 | 0.05 |
| Flavonoids | Scutellarein tetramethyl ether | R-111 | 0.59 | 0.22 | 0.11 | 0.14 | 0.36 |
|  |  | S-115 | 0.68 | 0.16 | 0.27 | 0.23 | 0.11 |
| Flavonoids | Quercetin-4'-O-glucuronide | R-111 | 0.95 | 0.41 | 0.17 | 0.41 | 0.26 |
|  |  | S-115 | 0.35 | 0.41 | 0.70 | 0.48 | 0.26 |
| Flavonoids | Quercetin-3-O-(6"-malonyl)galactoside | R-111 | 1.13 | 1.26 | 1.68 | 1.14 | 0.71 |
|  |  | S-115 | 0.00 | 0.04 | 0.06 | 0.08 | 0.16 |
| Flavonoids | Kaempferol-3,7-O-dirhamnoside (Kaempferitrin) | R-111 | 1.02 | 0.41 | 0.26 | 0.12 | 0.09 |
|  |  | S-115 | 1.39 | 0.63 | 0.52 | 0.31 | 0.20 |
| Flavonoids | Hispidulin-8-C-(2"-O-xylosyl)glucoside | R-111 | 1.85 | 4.54 | 4.27 | 4.59 | 4.21 |
|  |  | S-115 | 0.39 | 0.41 | 0.39 | 0.51 | 0.64 |
| Flavonoids | 4'-O-Glucosylvitexin | R-111 | 39.89 | 12.02 | 15.16 | 10.65 | 13.70 |
|  |  | S-115 | 6.30 | 5.68 | 5.38 | 5.27 | 2.90 |
| Flavonoids | Apigenin | R-111 | 0.00 | 0.09 | 0.08 | 0.08 | 0.07 |
|  |  | S-115 | 0.07 | 0.05 | 0.05 | 0.06 | 0.10 |
| Alkaloids | O-Phosphorylethanolamine | R-111 | 0.06 | 0.15 | 0.38 | 0.60 | 0.32 |
|  |  | S-115 | 0.41 | 0.70 | 1.17 | 1.29 | 1.26 |
| Alkaloids | Putrescine | R-111 | 0.00 | 7.69 | 6.79 | 9.56 | 7.69 |
|  |  | S-115 | 7.00 | 7.31 | 11.07 | 8.37 | 8.33 |
| Alkaloids | 5-Aminocycloheptane-1,2,3-triol | R-111 | 0.00 | 0.11 | 0.14 | 0.08 | 0.04 |
|  |  | S-115 | 0.16 | 0.15 | 0.22 | 0.07 | 0.10 |
| Alkaloids | 6,7-Dimethoxy-2-benzoxazolinone (DMBOA) | R-111 | 0.11 | 0.26 | 0.27 | 0.65 | 0.85 |
|  |  | S-115 | 0.14 | 0.21 | 0.25 | 0.26 | 0.46 |
| Alkaloids | Stearamide | R-111 | 0.56 | 0.46 | 0.44 | 0.46 | 2.48 |
|  |  | S-115 | 1.50 | 0.44 | 0.44 | 0.67 | 0.70 |
| Others | D-Glucose 1,6-bisphosphate | R-111 | 0.00 | 0.02 | 0.05 | 0.04 | 0.06 |
|  |  | S-115 | 0.02 | 0.05 | 0.09 | 0.08 | 0.04 |
| Others | Dihydroxyacetone phosphate | R-111 | 0.77 | 0.68 | 0.70 | 0.63 | 0.76 |
|  |  | S-115 | 0.19 | 0.61 | 1.00 | 1.18 | 0.82 |
| Others | 3-Methyl-1-pentanol | R-111 | 1.23 | 0.29 | 0.33 | 0.23 | 0.14 |
|  |  | S-115 | 0.28 | 0.28 | 0.06 | 0.07 | 0.09 |
| Others | 4-hydroxyphenyl acrylaldehyde | R-111 | 0.20 | 0.41 | 0.76 | 0.70 | 0.56 |
|  |  | S-115 | 0.39 | 0.64 | 0.46 | 0.78 | 0.54 |
| Others | 2-Amino-1,3-eicosanediol | R-111 | 13.76 | 6.51 | 4.52 | 2.00 | 1.21 |
|  |  | S-115 | 11.33 | 7.61 | 6.66 | 3.89 | 3.64 |
| Organic acids | Urocanic acid | R-111 | 0.00 | 0.25 | 0.22 | 0.30 | 0.48 |
|  |  | S-115 | 0.30 | 0.11 | 0.21 | 0.32 | 0.57 |
| Organic acids | 2,6-Diaminooimelic acid | R-111 | 1.46 | 0.51 | 0.52 | 0.25 | 0.14 |
|  |  | S-115 | 0.39 | 0.15 | 0.17 | 0.25 | 0.13 |
| Organic acids | D-Erythroneolactone | R-111 | 0.08 | 0.21 | 0.34 | 0.19 | 0.35 |
|  |  | S-115 | 0.17 | 0.27 | 0.37 | 0.17 | 0.20 |
| Amino acids derivatives | β-Alanine | R-111 | 0.09 | 0.05 | 0.08 | 0.09 | 0.14 |
|  |  | S-115 | 0.03 | 0.09 | 0.17 | 0.16 | 0.14 |
| Amino acids derivatives | Cyclo(Ala-Gly) | R-111 | 4.31 | 0.65 | 0.24 | 0.22 | 0.29 |
|  |  | S-115 | 2.24 | 0.84 | 0.22 | 0.13 | 0.16 |
| Amino acids derivatives | L-Lysine-Butanoic Acid | R-111 | 0.15 | 0.63 | 0.54 | 0.27 | 0.12 |
|  |  | S-115 | 0.69 | 1.69 | 2.29 | 2.93 | 2.84 |
| Lipids | 10-Undecenoic acid | R-111 | 0.02 | 0.06 | 0.07 | 0.05 | 0.06 |
|  |  | S-115 | 0.05 | 0.05 | 0.06 | 0.07 | 0.06 |
| Lipids | 1,18-Octadecanediol | R-111 | 0.01 | 0.03 | 0.04 | 0.03 | 0.01 |
|  |  | S-115 | 0.07 | 0.03 | 0.02 | 0.03 | 0.03 |
| Lipids | 12-Hydroxyoctadecanoic acid | R-111 | 0.00 | 0.01 | 0.37 | 0.14 | 0.26 |
|  |  | S-115 | 0.05 | 0.03 | 0.26 | 0.15 | 0.41 |
| Nucleotides derivatives | NADP (Nicotinamide adenine dinucleotide phosphate) | R-111 | 0.26 | 0.84 | 0.69 | 0.57 | 0.63 |
|  |  | S-115 | 0.22 | 0.48 | 1.26 | 0.70 | 1.07 |
| Quinones | 9,10-Dihydro-2-methoxy-4,5-phenanthrenediol | R-111 | 0.10 | 0.29 | 0.45 | 0.22 | 0.28 |
|  |  | S-115 | 0.29 | 0.35 | 0.20 | 0.21 | 0.32 |
| Lignans and Coumarins | Isolariciresinol | R-111 | 0.02 | 0.13 | 0.06 | 0.09 | 0.14 |
|  |  | S-115 | 0.07 | 0.07 | 0.00 | 0.10 | 0.15 |
| Terpenoids | 1β,2α,3α,19α,23-Pentahydroxyurs-12-en-28-oic acid | R-111 | 0.03 | 0.06 | 0.09 | 0.06 | 0.12 |
|  |  | S-115 | 0.02 | 0.05 | 0.04 | 0.07 | 0.10 |

\* Percentages of relative amounts.
† Mechanical damage and oral secretions treated for one hour (1 h).
‡ Mechanical damage and oral secretions treated for four hours (4 h).
§ Mechanical damage and oral secretions treated for eight hours (8 h).

Besides oral secretion with mechanical treatment, we also performed mechanical damage only, to find if mechanical damage can trigger the similar changes in metabolites. A total of 166 differential NOCs were found in resistant and susceptible lines, with 72 higher in resistant line and 94 higher in susceptible line. Differently, mechanical damage resulted in a decrease in the relative amounts of differentially expressed NOCs in sugarcane. In resistant line, 34.8 % of differential NOCs were inhibited. The NOCs in susceptible line were more affected by mechanical damage, with 54.7 % of differential NOCs inhibited. The inhibited differential NOCs in susceptible line were mainly flavonoids, amino acids derivatives and lipids, accounting for 15.1 %, 9.4 %, and 7.5 % of differential NOCs, respectively. There were also compounds that increased in amount in susceptible line, mainly phenolic acids, accounting for 9.4 % differential NOCs (**Table S8**). Therefore, we assumed that oral secretion plays more dominant role in triggering plant defense response than mechanical damage. Mechanical damage trends in shutting down plant metabolites.

### Effects of feeding with differential plant NOCs on *S. frugiperda*

In order to further explore those differential NOCs in their role in response to *S. frugiperda* damage, we selected two phenolic acids, one flavonoid and one lipid from 46 differential compounds. They are phenoxyacetic acid detected only in healthy resistant line, 4-methoxybenzaldehyde, apigenin and 12-hydroxyoctadecanoic acid detected only in healthy susceptible line (**Table 2**).

**Table 2.** Change of growth and development of *S*. *frugiperda* fed with diet supplemented with different selected NOCs and control diet.

| Compound information |  | the 3th- to 6th-instar larvae |  | pupa |  | Adult |  |
| --- | --- | --- | --- | --- | --- | --- | --- |
|  |  | Weight gain (g) | Period (days) | weight (g) | Period (days) | Pupation rate | Female/male |
| Phenolic acids | CK | 0.4104±0.0085 | 6.5294±0.1194 | 0.2120±0.0040 | 10.8000±0.0688 | 85.7% | 1 |
|  | 4-Methoxybenzaldehyde | 0.3900±0.0067 | 6.2571±0.0948 | 0.2148±0.0028 | 11.9643±0.0725 | 77.8% | 1.8 |
|  | Phenoxyacetic acid | 0.3866±0.0106 | 6.5556±0.1639 | 0.2112±0.0041 | 11.3200±0.1060 | 67.6% | 1.5 |
| Flavonoid | CK | 0.3783±0.0068 | 6.0500±0.1701 | 0.2061±0.0040 | 9.0909±0.1701 | 97.8% | 1.1 |
|  | Apigenin | 0.3404±0.0104 | 7.4105±0.2203 | 0.1765±0.0057* | 9.9402±0.2200 | 86.7% | 0.9 |
| Lipid | CK | 0.3016±0.0067 | 12.3105±0.0988 | 0.1923±0.0046 | 13.7011±0.0733 | 79.5% | 1 |
|  | 12-Hydroxyoctadecanoic acid | 0.3350±0.0070* | 10.7500±0.1609* | 0.1975±0.0040 | 12.0110±0.1609 | 85.8% | 0.9 |

The independence sample *t*-test are used, \*, $P < 0.05$ ; no “\*”, $P > 0.05$ .

Compared with the control diet, the body weight of the sixth-instar *S. frugiperda* larvae fed with diet supplemented with phenoxyacetic acid or 4-methoxybenzaldehyde decreased, with those with phenoxyacetic acid decreased the most (**Table 2**). The growth and development period of *S. frugiperda* larvae fed with 4-methoxybenzaldehyde was shortened, while those with phenoxyacetic acid was prolonged (**Table 2**). No significant effect on the pupal weight (*P* > 0.05), decrease in pupation rate and prolongation of the pupation period was observed (**Table 2**). In addition, these two phenolic acids supplement largely increased the male-to-female gender ratio of adult *S. frugiperda*, respectively, with 4-methoxybenzaldehyde supplement more significant (**Table 2**). It is noteworthy that both two phenolic acids contain benzene ring structure.

Compared with the control diet, the change of body weight from the third-instar *S. frugiperda* larvae to pupation with apigenin supplement decreased slightly with no significant difference (*P* > 0.05). The pupal weight was significantly decreased (*P* < 0.05), and the growth period of *S. frugiperda* was prolonged (**Table 2**). Change of body weight from the third-to sixth-instar *S. frugiperda* larvae with 12-hydroxyoctadecanoic acid supplement increased significantly by 11.7 %, (*P* < 0.05). The developmental period was shortened significantly by 12.74 % (*P* < 0.05). In addition, the pupal weight and pupation rate of *S. frugiperda* fed with 12-hydroxyoctadecanoic acid increased with no significant difference (*P* > 0.05). The male-to-female ratio of *S. frugiperda* fed with 12-hydroxyoctadecanoic acid had no significantly difference (*P* > 0.05) (**Table 2**).

### Differences in plant VOCs between resistant and susceptible sugarcane lines

In the healthy state and under insect damage, plant VOCs may be volatized out of plants as identification signals that affect the host selection of herbivore insects. In this study, VOCs were measured using HS-SPME at 0h, 1h, 4h, and 8h time points before/after *S. frugiperda* treatment (**Table S9** and **Figure S4**). We selected 102 plant-derived VOCs with match factors above 60 for subsequent analyses, excluding the non-plant-derived VOCs (**Table S10**). Among the 102 VOCs, 18 plant VOCs were higher in relative amounts before/after treatment in susceptible line and 73 plant VOCs were lower in susceptible line (**Figure 2A,B,C**). The other 11 plant VOCs showed lower relative amounts in healthy state and 4 h treatment in susceptible line and higher amount at 8 h treatment (**Figure 2A**).

**Figure 2.**
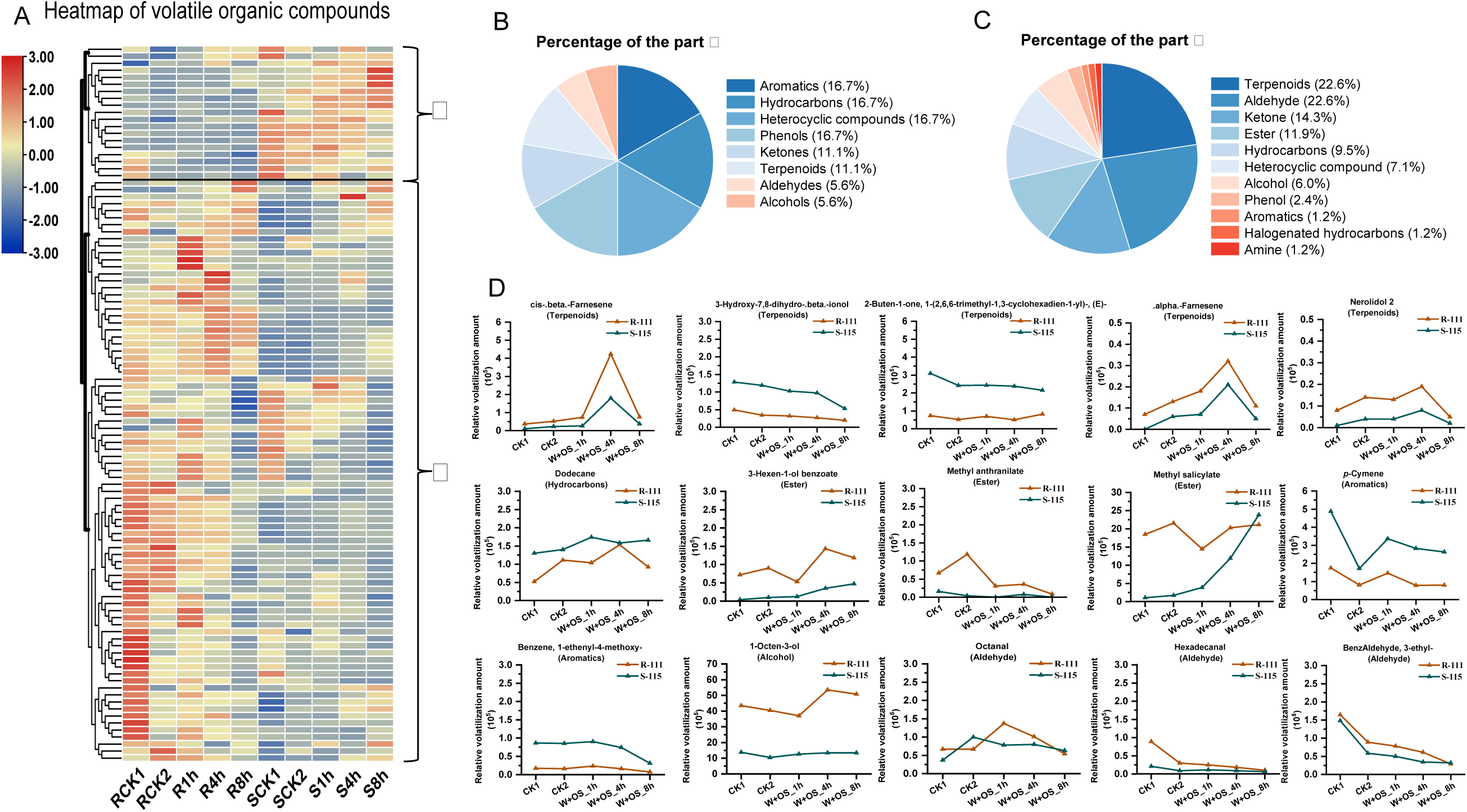
Analysis of VOCs in resistant and susceptible sugarcane lines. (A) Heatmap of 102 VOCs levels in different treatment of resistant and susceptible sugarcane lines. (B) Category proportion of 18 compounds higher in susceptible line. (C) Category proportion of 83 compounds higher in resistant line. (D) Line graph of important compounds levels. R-111: medium *S*. *frugiperda*-resistant lines; S-115: highly *S. frugiperda*-susceptible lines.

In the healthy state, the largest proportion of plant VOCs of resistant line were terpenoids, accounting for 20.59 %, while in the susceptible line were aldehydes, accounting for 19.8 % (**Table S11**). Oral secretions of *S. frugiperda* combined with mechanical damage triggered differential plant VOC changes in both the resistant and susceptible lines. In resistant line, level of aldehydes increased rapidly after treatment constituting the largest proportion for about 36.4 % and 26.7% at 1 h and 4h respectively, followed by heterocyclic compounds (**Table S12**). At 8 h treatment, the proportion of aromatics increased, accounting for approximately 9.1 %, which was similar to terpenoids and heterocyclic compounds. In susceptible line, the amounts of esters and terpenoids rapidly increased following treatment. With an increase in treatment time, the amounts of ketones, aldehydes, and heterocyclic compounds increased. At 1 h treatment, terpenoids constituted the largest proportion for 35.7 %, followed by esters. At 4 h, the amounts of aldehydes increased rapidly to the largest, accounting for 20.8 %. At 8 h, terpenoids still accounted for the largest proportion, for 28.1 %, followed by aldehydes and esters (**Table S13**). The rapid change of plant VOC contents reflects the defense process of plants in response to insect infestation. Therefore, we analyzed the changes of plant VOCs and divided them into 12 types of trends (**Figure S5**).

Mechanical damage without oral secretion resulted in a decrease in VOCs similarly with NOCs. In the resistant line, the relative amount of plant VOCs reduced, except for hydrocarbons (dodecane) (**Table S15**). In contrast, the plant VOCs in susceptible line were less affected by mechanical damage, with only 66.7 % of differential plant VOCs inhibited. We speculate that this inhibition caused by mechanical damage might be due to the protective mechanism of sugarcane against further damage.

### Identification of plant VOCs attractive and repellent to *S. frugiperda*

We further screened individual compounds to test whether they might attract or repel *S. frugiperda*. Among the 102 plant VOCs, *α*-farnesene was detected in the healthy resistant line only (**Table S10**). *α*-farnesene functions to defend plants from insects (Huang et al., 2022). Therefore, we speculated that it also has a repellent effect on *S. frugiperda*. The relative amounts of two aromatics, benzene, 1-ethenyl-4-methoxy– and *p*-cymene, were 5.1 and 2.8 times higher, respectively, in healthy susceptible line than those in resistant line. The relative amount of dodecane was 2.5 times higher in susceptible line than in resistant line (**Table S11**). Methyl jasmonate, a major insect-defense compound relating to the jasmonic acid pathway, was not detected in either resistant or susceptible lines. However, the relative amount of methyl salicylate, relating to the SA pathway, was lower in healthy susceptible line than in resistant line (**Table S11**). When comparing different time points before and after infestation, 1-octen-3-ol and methyl anthranilate increased rapidly in susceptible line, although the relative amounts were still lower than those in the resistant line (**Tables S12-S14**). We speculated that 1-octen-3-ol and methyl anthranilate may function as repellents for *S. frugiperda*. In addition, dodecane was upregulated after treatment in both resistant and susceptible lines. Its relative amount in susceptible line was higher than that in resistant line, which may be attractive to *S. frugiperda*. Most of these plant VOCs, especially dodecane, were downregulated after mechanical damage (**Table S15**). This overall reduction indicates that mechanical damage may inhibit the production of plant VOCs. Finally, 15 compounds with differences were screened. These compounds were assumed potentially attractive or repellent to *S. frugiperda* and may affect the preferences of *S. frugiperda* (**Figure 2D** and **Table 3**).

**Table 3.**
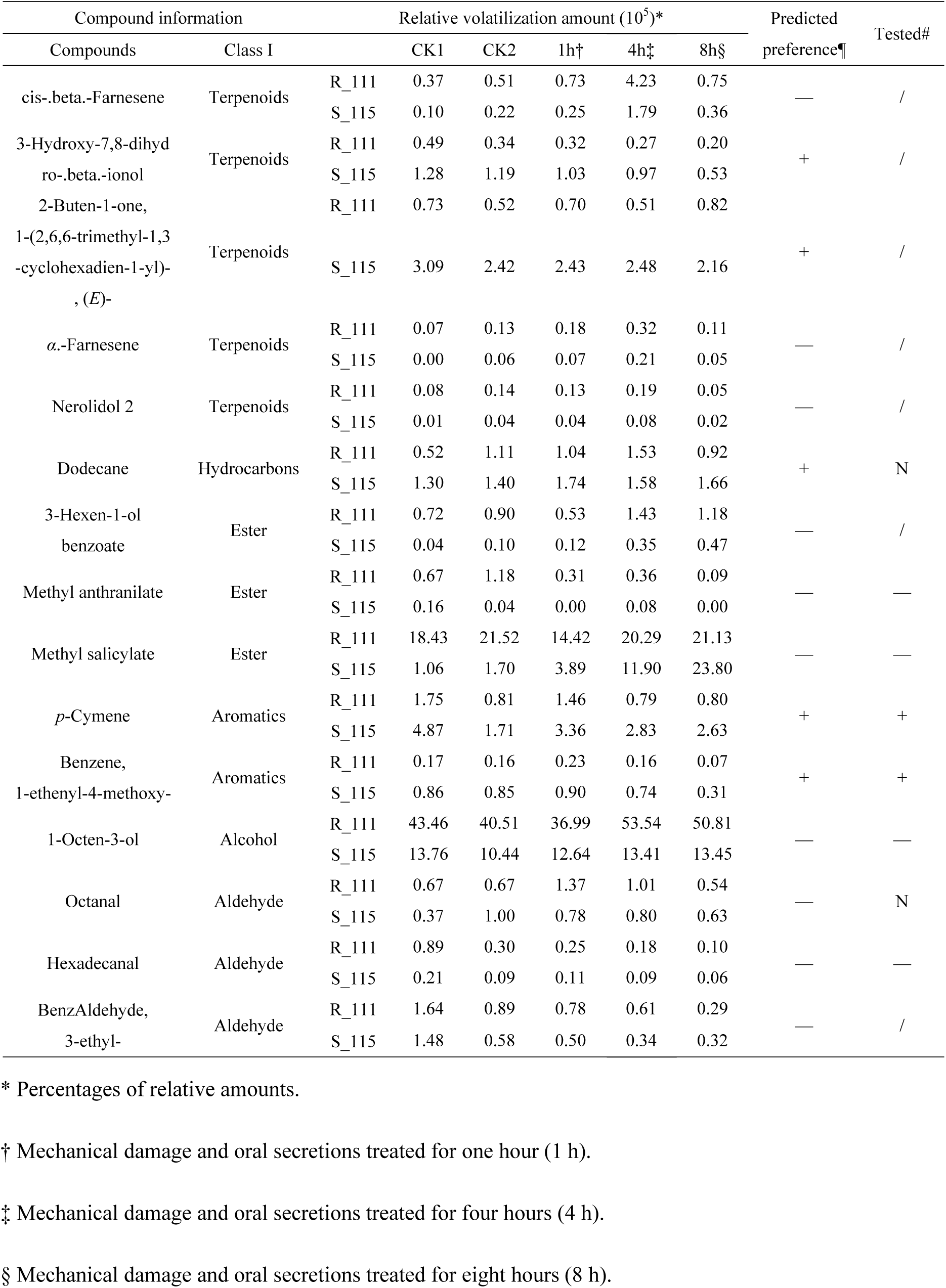
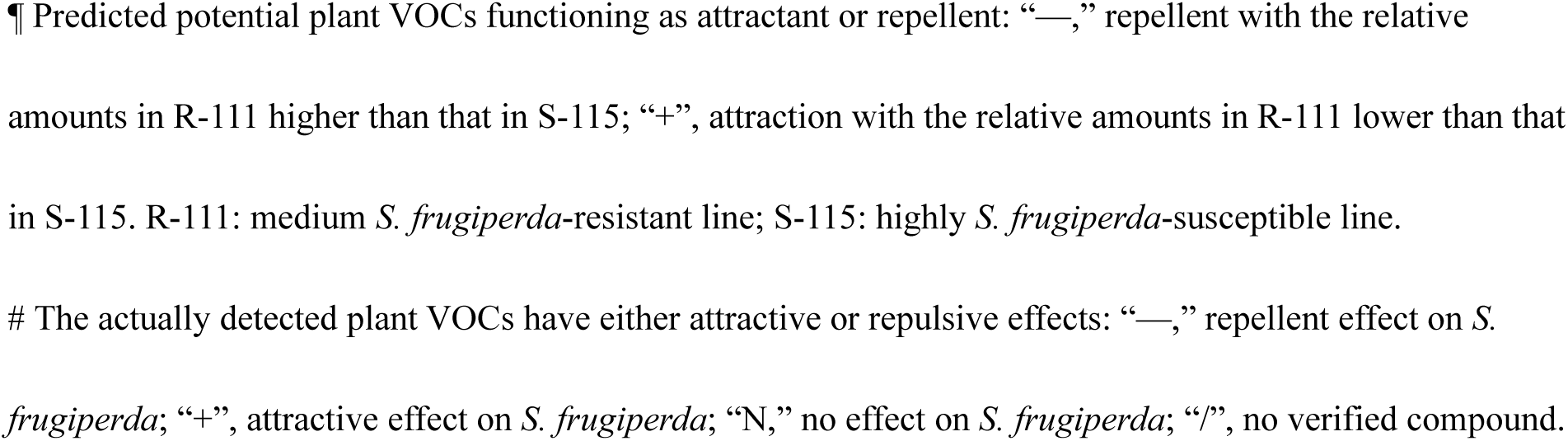
Information on selected compounds based on plant VOC analysis.

| Compound information |  | Relative volatilization amount (10<sup>5</sup>)* |  |  |  |  |  | Predicted preference¶ | Tested# |
| --- | --- | --- | --- | --- | --- | --- | --- | --- | --- |
| Compounds | Class I |  | CK1 | CK2 | 1h† | 4h‡ | 8h§ |  |  |
| cis-.beta.-Farnesene | Terpenoids | R_111 | 0.37 | 0.51 | 0.73 | 4.23 | 0.75 | — | / |
|  |  | S_115 | 0.10 | 0.22 | 0.25 | 1.79 | 0.36 |  |  |
| 3-Hydroxy-7,8-dihydro-.beta.-ionol | Terpenoids | R_111 | 0.49 | 0.34 | 0.32 | 0.27 | 0.20 | + | / |
|  |  | S_115 | 1.28 | 1.19 | 1.03 | 0.97 | 0.53 |  |  |
| 2-Buten-1-one, 1-(2,6,6-trimethyl-1,3-cyclohexadien-1-yl)-, (E)- | Terpenoids | R_111 | 0.73 | 0.52 | 0.70 | 0.51 | 0.82 | + | / |
|  |  | S_115 | 3.09 | 2.42 | 2.43 | 2.48 | 2.16 |  |  |
| α-Farnesene | Terpenoids | R_111 | 0.07 | 0.13 | 0.18 | 0.32 | 0.11 | — | / |
|  |  | S_115 | 0.00 | 0.06 | 0.07 | 0.21 | 0.05 |  |  |
| Nerolidol 2 | Terpenoids | R_111 | 0.08 | 0.14 | 0.13 | 0.19 | 0.05 | — | / |
|  |  | S_115 | 0.01 | 0.04 | 0.04 | 0.08 | 0.02 |  |  |
| Dodecane | Hydrocarbons | R_111 | 0.52 | 1.11 | 1.04 | 1.53 | 0.92 | + | N |
|  |  | S_115 | 1.30 | 1.40 | 1.74 | 1.58 | 1.66 |  |  |
| 3-Hexen-1-ol benzoate | Ester | R_111 | 0.72 | 0.90 | 0.53 | 1.43 | 1.18 | — | / |
|  |  | S_115 | 0.04 | 0.10 | 0.12 | 0.35 | 0.47 |  |  |
| Methyl anthranilate | Ester | R_111 | 0.67 | 1.18 | 0.31 | 0.36 | 0.09 | — | — |
|  |  | S_115 | 0.16 | 0.04 | 0.00 | 0.08 | 0.00 |  |  |
| Methyl salicylate | Ester | R_111 | 18.43 | 21.52 | 14.42 | 20.29 | 21.13 | — | — |
|  |  | S_115 | 1.06 | 1.70 | 3.89 | 11.90 | 23.80 |  |  |
| p-Cymene | Aromatics | R_111 | 1.75 | 0.81 | 1.46 | 0.79 | 0.80 | + | + |
|  |  | S_115 | 4.87 | 1.71 | 3.36 | 2.83 | 2.63 |  |  |
| Benzene, 1-ethenyl-4-methoxy- | Aromatics | R_111 | 0.17 | 0.16 | 0.23 | 0.16 | 0.07 | + | + |
|  |  | S_115 | 0.86 | 0.85 | 0.90 | 0.74 | 0.31 |  |  |
| 1-Octen-3-ol | Alcohol | R_111 | 43.46 | 40.51 | 36.99 | 53.54 | 50.81 | — | — |
|  |  | S_115 | 13.76 | 10.44 | 12.64 | 13.41 | 13.45 |  |  |
| Octanal | Aldehyde | R_111 | 0.67 | 0.67 | 1.37 | 1.01 | 0.54 | — | N |
|  |  | S_115 | 0.37 | 1.00 | 0.78 | 0.80 | 0.63 |  |  |
| Hexadecanal | Aldehyde | R_111 | 0.89 | 0.30 | 0.25 | 0.18 | 0.10 | — | — |
|  |  | S_115 | 0.21 | 0.09 | 0.11 | 0.09 | 0.06 |  |  |
| Benzaldehyde, 3-ethyl- | Aldehyde | R_111 | 1.64 | 0.89 | 0.78 | 0.61 | 0.29 | — | / |
|  |  | S_115 | 1.48 | 0.58 | 0.50 | 0.34 | 0.32 |  |  |

\* Percentages of relative amounts.
† Mechanical damage and oral secretions treated for one hour (1 h).
‡ Mechanical damage and oral secretions treated for four hours (4 h).
§ Mechanical damage and oral secretions treated for eight hours (8 h).
¶ Predicted potential plant VOCs functioning as attractant or repellent: “—,” repellent with the relative amounts in R-111 higher than that in S-115; “+”, attraction with the relative amounts in R-111 lower than that in S-115. R-111: medium *S. frugiperda*-resistant line; S-115: highly *S. frugiperda*-susceptible line.

To further investigate the effects of plant VOCs on *S. frugiperda*, we conducted insect behavioral preference assays on *S. frugiperda* larvae, adult females, and adult males. We tested eight selected plant VOCs, excluding those that were unavailable (no information on CAS number or commercially unavailable) (**Table 3**). Two aromatic compounds, benzene, 1-ethenyl-4-methoxy– and *p*-cymene, were attractive to *S. frugiperda*. Benzene, 1-ethenyl-4-methoxy-, had a significant (*P* < 0.05) attraction effect on adult females and larvae, but not on adult males. *p*-Cymene attracted larvae, adult males and females of *S. frugiperda*. Dodecane was predicted to be an attractive compound (**Table 3**); however, it had no significant (*P* > 0.05) effect on *S. frugiperda.* Methyl anthranilate and 1-octen-3-ol were repellent to larvae, adult males and females of *S. frugiperda*. Methyl salicylate was significantly (*P* < 0.05) repellent to *S. frugiperda* adult females and larvae, but not to adult males. Hexadecanal only had a repellent effect on *S. frugiperda* adult males. Octanal was previously predicted to be a repellent compound to *S. frugiperda* but had no significant (*P* > 0.05) effect on *S. frugiperda* preferences (**Figure 3**). Based on the results of the insect behavioral preference assays, we found that most of the plant VOCs with higher relative amounts in resistant line had repellent effects on *S. frugiperda*, whereas the two aromatics with higher relative amounts in the susceptible line were attractive to *S. frugiperda*. Notably, both of these aromatics contain benzene ring structure.

**Figure 3.**
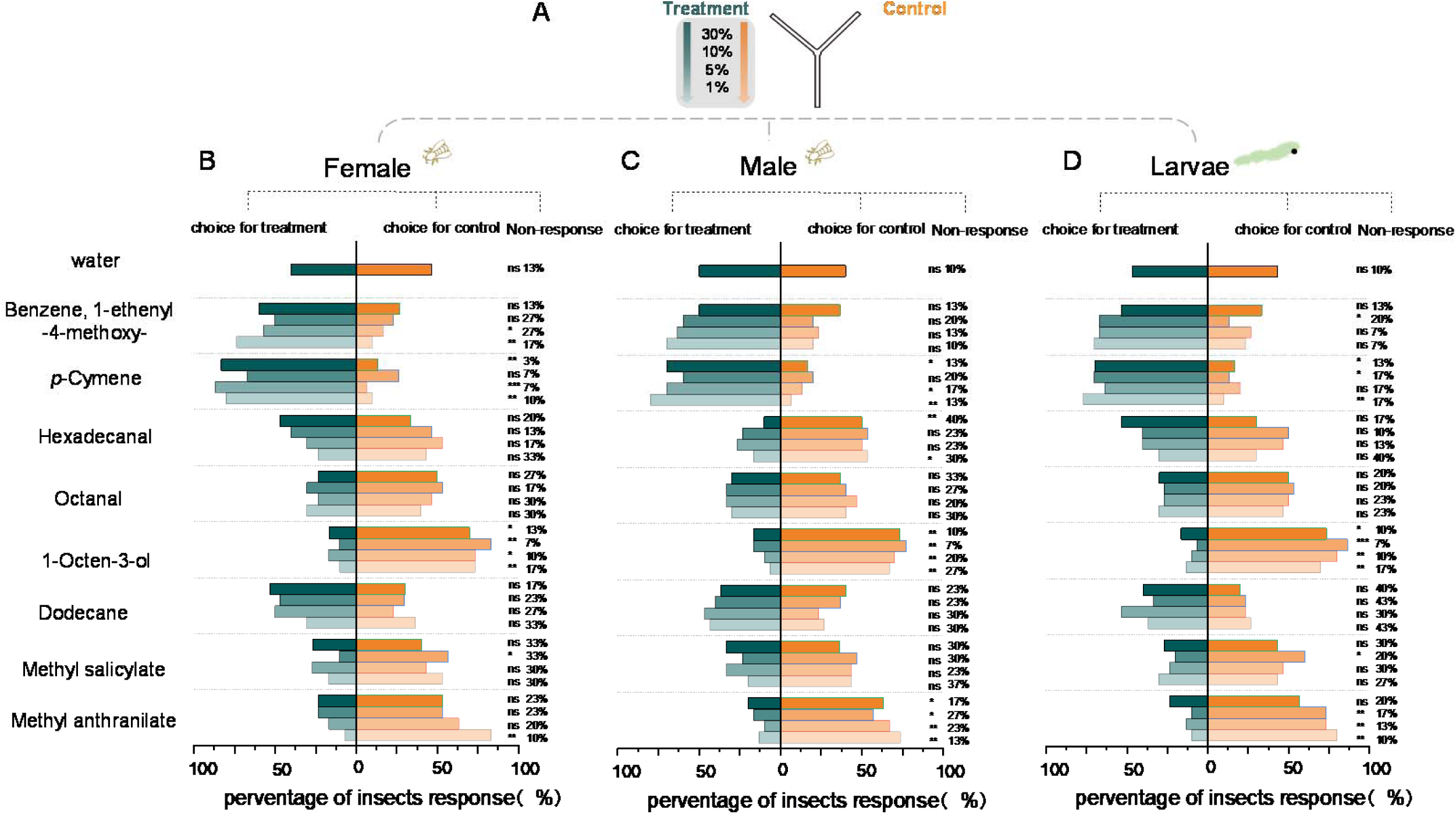
Insect olfactory behavioral preference test. (A) Compounds of different concentrations with water as control were tested for insect response using Y-tube. (B) Adult females, (C) adult males and (D) larvae of *S. frugiperda* choice with different concentrations of volatile compound. The asterisks are based on Chi-Squared test, ***, *P* < 0.001; **, *P* < 0.01; *, *P* < 0.05; ns, *P* > 0.05.

### Tryptophan, an aromatic compound precursor containing benzene ring, could promote the growth and development of *S. frugiperda*

Base on the above results, we noted that benzene ring containing compounds in both plant NOCs and VOCs might play important role in resistance or susceptibility of plant to attack of *S. frugiperda*. Therefore, we further analyzed the transcriptome of the third-instar *S. frugiperda* larvae fed on the resistant and susceptible sugarcane leaves, in order to find out if relevant pathway or genes can be screened. The differential genes were used to perform KEGG analysis (**Figure 4A**). Pathways identified related to phenylalanine, tyrosine and tryptophan biosynthesis, phenylalanine metabolism, folate biosynthesis, biosynthesis of amino acids, and disease related process (**Figure 4B**). All phenylalanine, tyrosine, tryptophan and folate contain benzene ring. The differential gene *phhA*, encoding phenylalanine 4-monooxygenase, and *SLC39A5* relating to disease were annotated. As in this study, aromatic phenolic compounds from susceptible line were found to affect *S. frugiperda* growth and development and two aromatic VOCs from susceptible line can attract *S. frugiperda*, we further analyzed the precursor of aromatics, tryptophan, which was identified in KEGG as well. Tryptophan is closely related to the synthesis of downstream aromatic compounds (Barik, 2020). Artificial diets supplemented with tryptophan were used to determine the changes in body weight, pupation rate, and developmental period of third– to sixth-instar *S. frugiperda* larvae (**Figure 4C**, **D**, **E**). The average body weight of *S. frugiperda* fed with the control diet, diet supplemented with half of maize powder, diet supplemented with tryptophan was 0.378 g, 0.364 g, and t 0.341 g, respectively (**Figure 4E**). The results showed that feeding with diet supplemented with tryptophan reduced the body weight of *S. frugiperda,* but with no significance (*P* > 0.05). Compared with diet supplemented with half of maize powder, the development period of the third-to-sixth-instar *S. frugiperda* larvae fed with half of maize powder supplemented with tryptophan increased from six days to eight days, and the pupation rate increased from 6.67 % to 26.67 % (**Figure 4D**). Hence, tryptophan was beneficial to *S. frugiperda* development in nutritional deficiency condition.

**Figure 4.**
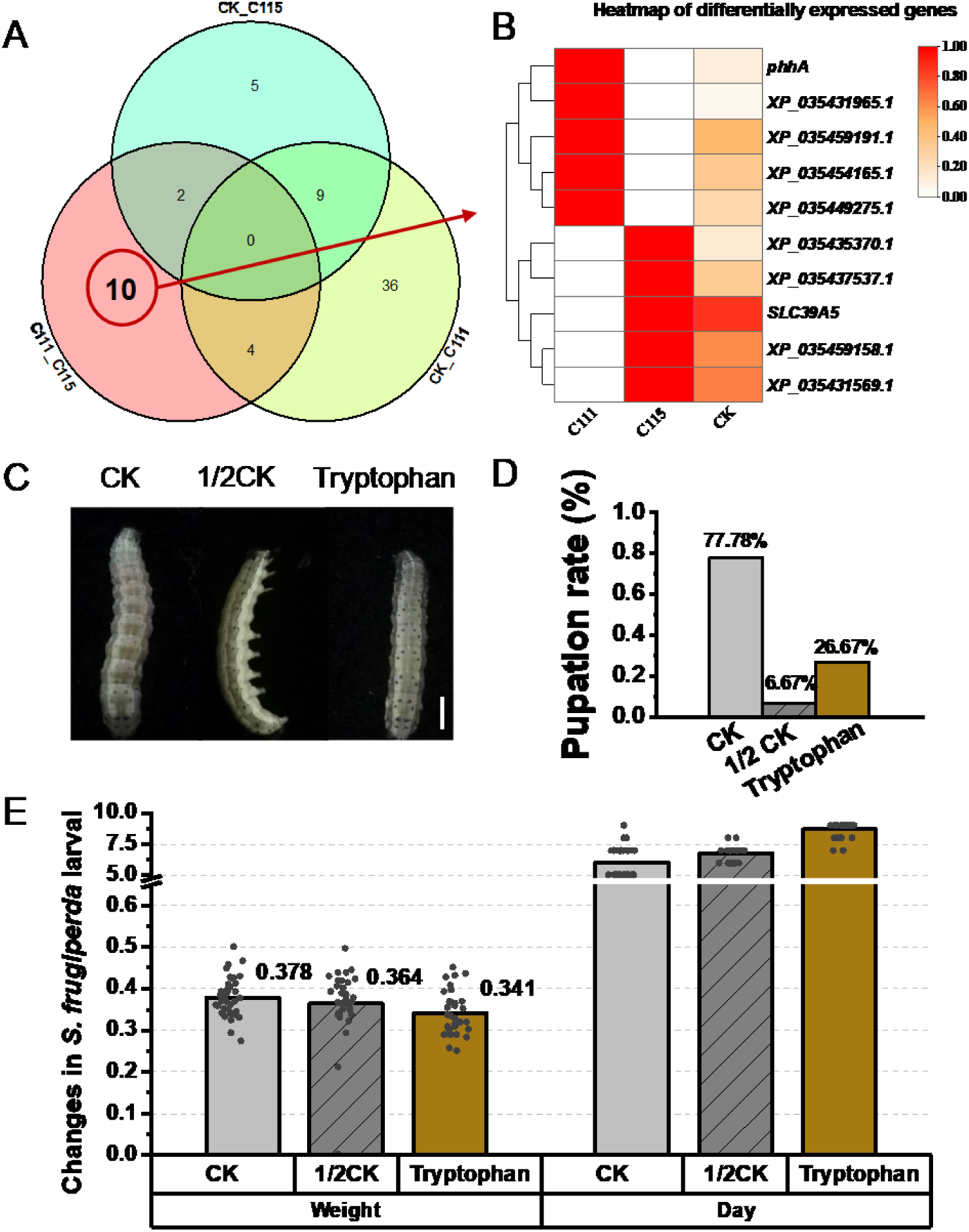
Transcriptome analysis revealed that effects of tryptophan on the growth and development of *S. frugiperda*. (A) Venn diagram of differential genes in different comparison groups of *S. frugiperda*. (B) heatmap of differential gene expression levels. C111: *S*. *frugiperda* fed on the resistant line, C115: *S. frugiperda* fed on the susceptible line. (C) The sixth instar *S. frugiperda* larvae fed with different artificial diets, bars, 1 cm. (D) Pupation rate of *S. frugiperda* fed with different artificial diets. (E) Body weight gain and growth cycle of *S. frugiperda* fed with different artificial diets.

## Conclusion

In summary, we find in sugarcane NOCs, VOCs and the specific aromatic precursor that benzene ring containing compounds play critical role in plant-*S. frugiperda* interaction. Phenolic acids containing benzene ring accounted for the largest proportion of differential NOCs. Levels of 66.7% of these phenolic acids were higher in susceptible line. The phenolic acids in sugarcane leaves could improve the male-to-female ratio of adult *S. frugiperda*. Aromatics containing benzene ring, accounted for the largest of differential VOCs in susceptible line. The volatile aromatics from sugarcane leaves could attract *S. frugiperda* and influence its selection of host plants. Tryptophan in sugarcane leaves that related to the synthesis of downstream aromatic compounds was conducive to improving the pupation rate of *S. frugiperda* in nutritional deficiency condition. Therefore, we suggested that the compounds with benzene ring structure especially phenolic acids and aromatics were important factors affecting the interaction between sugarcane and *S. frugiperda* (**Figure 5**).

**Figure 5.**
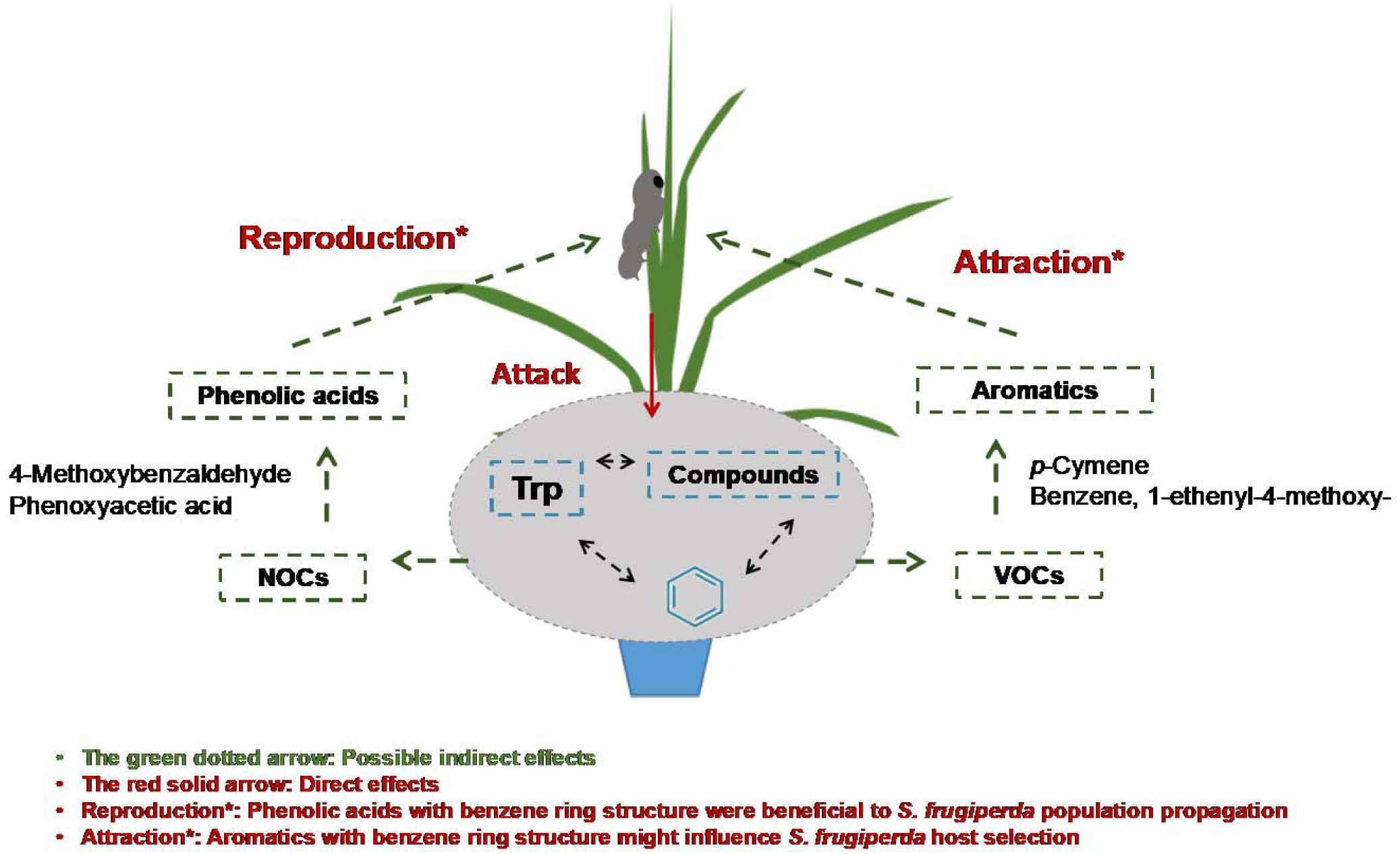
The aromatic compounds with benzene ring structure in resistant and susceptible sugarcane lines were important factors affecting the interaction between sugarcane and *S. frugiperda*. The tryptophan, as an aromatic amino acid, is the upstream compound of aromatic compounds, and may positively correlated with the synthesis of aromatic compounds The phenolic acids could increase the male-to-female ratio of *S. frugiperda*, and aromatic volatiles had an attractive effect on *S. frugiperda*. These compounds contain a benzene ring structure.

## Discussion

Plant secondary metabolites are one of the factors that affect the interaction between plants and herbivorous insects and are important immediate responses of plants to herbivorous insect infestation (War et al., 2011). In this study, we found differences in plant secondary metabolites between resistant and susceptible sugarcane lines from the same genetic population in the healthy state and before and after *S. frugiperda* infestation, and identified some important NOCs affecting the growth and development of *S. frugiperda*, and some important VOCs with attractive or repellent effects on *S. frugiperda*.

Plants can resist insect predation by synthesizing a range of active small molecule compounds (secondary metabolites). The main plant secondary metabolites can be divided into phenolic acids, terpenoids and nitrogen compounds. These secondary metabolites participate in plant defense and affect the growth and development of insects (Tao et al., 2012; Erb and Kliebenstein, 2020). Insects use secondary metabolites as informational chemical compounds and try to avoid the negative effects of secondary metabolites through choice behavior and metabolic adaptations while protecting themselves from predator insects (Nishida, 2002; Behmer, 2009; Opitz and Müller, 2009; Stahl et al., 2018). It has been reported that whether insects can effectively utilize ingested phenolic acids depends on its antioxidants, such as cytochrome P450 (Rey et al., 1999; Simmonds, 2003). Insects with excessive intake of phenolic acids or lack of related antioxidants are unable to use these compounds, negatively affecting their own growth and development. Insects avoid these situations during host selection (Simmonds, 2003). Different from these conclusions, the relative amounts of phenolic acids in the susceptible line were higher than that in the resistant line. In order to further explore the reasons for the opposite conclusion, the effects of phenolic acids on the growth and development of *S. frugiperda* were studied by feeding artificial diets supplemented with phenolic acids. The results showed that phenolic acids 4-methoxybenzaldehyde and phenoxyacetic acid decreased the body weight and pupation rate of *S. frugiperda.* Interestingly, these two phenolic acids both increased the male-to-female ratio of *S. frugiperda*. As an invasive pest, the male-to-female ratio of *S. frugiperda* is an important basis for emigrant and immigrant population, and the increasing male-to-female ratio may be conducive to emigrant and immigrant population of *S. frugiperda* (Riley et al., 2009). The activity of oxidase and hydrolase in *S. frugiperda* is higher than that of other insects, which is conducive to the synthesis of antioxidants and effective utilization of phenolic acids (Gouin et al., 2017; Silva-Brandão et al., 2017). Based on these researches, we speculated that the high relative amounts of phenolic acids in the susceptible line was beneficial to the breeding of *S. frugiperda*, and *S. frugiperda* was more inclined to choose the susceptible line as the host plant.

Flavonoids as plant component are widely found in nature. Flavonoids are often used as antioxidants and free radical scavengers to enhance the immune regulatory capacity of insects (Williams et al., 2004). Apigenin, as an antioxidant among flavonoids, can directly eliminate or reduce the level of intracellular reactive oxygen species or reduce the content of malondialdehyde (the end product of lipid oxidation). It can improve the enzyme activities of antioxidant such as SOD, CAT and GSH, to protect and maintain normal growth of insects (Hu et al., 2016; Han et al., 2017). There was no significantly (*P* > 0.05) effect of apigenin on the growth and development of *S. frugiperda*. Lipids in plants are energy sources for insect reproduction, embryonic development and metamorphosis. They play an important role in regulating insect activity and conveying insect information (Kameoka and Gutjahr, 2022). It is similar to our findings that lipids in sugarcane leaf can significantly increase the weight of *S. frugiperda* larvae and shorten the development period of *S. frugiperda*. Further, lipids in sugarcane leaf also increased the pupal weight and pupation rate of *S. frugiperda*. After the insect infestation, the secondary metabolites in sugarcane synthesize volatile secondary metabolites through a series of pathways, which further become the recognition signal of insect behavior selection (Erb and Kliebenstein, 2020).

Different plant VOCs have different effects on insect preferences (Hu, 2022). This study investigated the effects of different plant VOCs on the preferences of *S. frugiperda*. Previous studies have shown that terpenoids are important HIPVs in direct or indirect plant defense, as they not only repel herbivorous insects and also attract predators or parasites of herbivorous insects (Heil, 2008; War et al., 2011; Sharma et al., 2017; Huang et al., 2022). Furthermore, heterocyclic compounds have been reported to be toxic to the larvae of *S. frugiperda* (Huang et al., 2019; Kim et al., 2020). These results are consistent with our findings which show that terpenoids and heterocyclic compounds are more abundant in the resistant line than in the susceptible line. Aldehydes are HIPVs that are generally regarded as natural plant insecticides and are effective eco-chemical means of pest control (Hubert et al., 2008). The aldehyde compound hexadecanal is mostly used to investigate the interaction of insect females and males; however, few studies have explored the interaction between hexadecanal and insects (Choi et al., 2016). We found that hexadecanal had a weak repellent effect on adult male *S. frugiperda*. Octanal, an aldehyde volatilized from damaged plants, is considered to be a weak insect repellent (Laznik and Trdan, 2016). However, we did not observe any effect of octanal on the preferences of *S. frugiperda* in this study. The ester compound methyl anthranilate was found in other studies to have a repellent effect on insects. Higher concentrations of methyl anthranilate can repel and inhibit the adult emergence of *Drosophila suzukii* (Bräcker et al., 2020; Vuts et al., 2021). This is consistent with our findings which show that methyl anthranilate has a repellent effect on *S. frugiperda*. Notably, the alcohol 1-octen-3-ol, can be rapidly synthesized and volatilized after mechanical damage and has a repellent effect on the southern house mosquito (Xu et al., 2015; Ntoruru et al., 2022), which is also consistent with our results. Some aromatics have been reported to exert attractive effects on other insects. 4-Ethylguaiacol and 4-methylphenol are aromatics that attract Drosophila (Brown et al., 2017). Aromatic *p*-cymene is attractive to *Leptocybe invasa* and benzene, 1-ethenyl-4-methoxy– to *Erioscelis emarginata* beetles (Dötterl et al., 2012; Huang et al., 2022). The results of these previous studies are consistent with the observed effects of these plant VOCs on *S. frugiperda*. Aromatics play an important role in the interdependence of plants and insects. Phenolic acids are a kind of aromatics, which are usually used as the precursor of synthesis of aromatic VOCs. The results showed that the aromatics had an attractive effect on *S. frugiperda* and was conducive to the population reproduction of *S. frugiperda*. We speculate that the aromatics in resistant and susceptible sugarcane lines play an important role in the sugarcane-*S. frugiperda* interaction mechanisms.

The research showed that tryptophan, as an aromatic amino acid, is the upstream compound of aromatic compounds, and may positively correlated with the synthesis of aromatic compounds. It plays an important role in the regulatory network of plant defense response (Barik, 2020). For insects, tryptophan can help maintain the normal physiological functions of insects under stress (Gao et al., 2022). These previous results are consistent with our finding that the expression level of *phhA* related to tryptophan synthesis pathway in *S. frugiperda* fed with resistant line was higher than that fed with susceptible line, and tryptophan is required for the pupation of *S. frugiperda* in unfavorable conditions. Therefore, *S. frugiperda* may prefer susceptible line as hosts because they release more aromatics and produce more tryptophan compared to resistant line. Based on the results, we hypothesize that the differences in aromatic compounds between resistant and susceptible lines may be related to *S. frugiperda* host selection. The reasons for the differences in aromatic compounds between resistant and susceptible sugarcane lines is worth of further investigation.

## Materials and methods

### Plant materials and insect oral secretion collection

The plant material was sugarcane line S-115 that was found to be highly susceptible to *S. frugiperda* in two-year field evaluations and resistance identification. It was obtained from a genetic population using Hocp00-1142 and Yacheng 06-92 as parents. R-111 (medium resistance) is a sister line obtained from the same genetic population. The plants were planted in the Germplasm Resource Nursery of Guangxi University, located in Fusui County, Guangxi Province, China. Similar healthy leaves from 3 replicative plants in the late seedling stage from each line were selected as the experimental materials. *S. frugiperda* was collected from the experimental field. Laboratory rearing and subculturing were made on 9 cm Petri dishes supplemented with maize leaves and maize seeds. The oral secretions of third-instar larvae incubated by *S. frugiperda* eggs produced from artificial rearing were collected in empty 1.5 mL centrifuge tubes. The oral secretion collection method was performed according to Si (Si et al., 2020). The oral secretion samples in centrifugal tube was placed on ice and stored at –80℃.

### Treatments

Insect oral secretions were mixed well before use. The positive third leaves of sugarcane plants planted in the field were selected as the experimental leaves. The mechanical damage method was performed as described by Si (Si et al., 2020). Scissors were used to mimic the *S. frugiperda* bite damage. The 20 μL of oral secretions were placed at the mechanical wounds for treatment. Five time-points were set for this experiment: untreated (CK1), mechanical damage for one hour (CK2), mechanical damage and oral secretion treatment for one hour (W+OS_1h), mechanical damage and oral secretion treatment for four hours (W+OS_4h), and mechanical damage and oral secretion treatment for eight hours (W+OS_8h). The treatment and collection timess were strictly controlled. Sugarcane leaves for each test, conducted in triplicate, were rapidly frozen in liquid nitrogen and stored at –80℃.

### *S. frugiperda* resistance evaluation

Four healthy sugarcane plants, R-111 and S-115, with similar growth status at the seedling stage, were planted in a greenhouse. Third-instar *S. frugiperda* for artificial rearing was placed on the first positive leaves of sugarcane. Insect growth and development were recorded daily. Inoculation ceased at the end of the third instar stage. Plant status was recorded and evaluated according to the degree of damage. The damage degree was divided into 5 levels: “A,” slightly damaged or no trace of infestation; “B,” little trace of infestation; “C,” growing point was mildly damaged; “D,” growing point was broken but didn’t affect plant growth; and “E,” growing point was broken and plant growth was affected.

### Secondary metabolites analysis

Leaf samples were collected from resistant and susceptible sugarcane lines at different time points before and after insect infestation. Leaf samples were sent to Metware Biotechnology Co., Ltd. for NOC and VOC measurements. The NOC analysis method was performed based on Wang (Wang et al., 2021). The sample extracts were analyzed using an UPLC-ESI-MS/MS system (UPLC, SHIMADZU Nexera X2; MS, Applied Biosystems 4500 Q TRAP). The analytical conditions were as follows, UPLC: column, Agilent SB-C18 (1.8 µm, 2.1 mm x 100 mm); The mobile phase was consisted of solvent A, pure water with 0.1 % formic acid, and solvent B, acetonitrile with 0.1 % formic acid. Sample measurements were performed with a gradient program that employed the starting conditions of 95 % A, 5 % B. Within 9 min, a linear gradient to 5 % A, 95 % B was programmed, and a composition of 5 % A, 95 % B was kept for 1 min. Subsequently, a composition of 95 % A, 5.0 % B was adjusted within 1.1 min and kept for 2.9 min. The flow velocity was set as 0.35 mL per minute; The column oven was set to 40°C; The injection volume was 4 μL. The effluent was alternatively connected to an ESI-triple quadrupole-linear ion trap (QTRAP)-MS.

The VOC analysis method was performed based on Huang (Huang et al., 2019). Samples were transferred into a 20 mL head-space vial (Agilent, Palo Alto, CA, USA) containing an NaCl saturated solution to inhibit any enzyme reaction. Fully automatic headspace-solid phase microextraction (HS-SPME) was used for sample extraction for gas chromatography-mass spectrometry (GC-MS) analysis. At the time of SPME analysis, each vial was placed at 100°C for 5 min. Thereafter, a 120 µm divinylbenzene/carboxen/polydimethylsilioxan fiber (Agilent) was exposed to the headspace of the sample for 15 min at 100°C. VOCs were identified and quantified using an Agilent Model 8890 GC and 5977 B mass spectrometer (Agilent). Mass spectra were scanned in the range of m/z 50–500 amu at 1 s intervals. The software Qualitative Analysis Workflows B.08.00 was used to view the raw data from the machine and conduct mass spectrometry and qualitative analyses. Data dispersion was analyzed using an Empirical Cumulative Distribution Function (ECDF).

The secondary metabolites were identified by comparing the mass spectra with the data system library (MWGC or NIST) and the linear retention index. PubChem (https://pubchem.ncbi.nlm.nih.gov/) was used to identify the secondary metabolites sources. Principal component analysis (PCA) was performed using the statistical function prcomp within R. Pearson correlation coefficients (PCCs) between samples were calculated using the cor function in R (v4.1.2) and are presented as heatmaps. Secondary metabolites with significant differences in content, indicated by a fold change ≥ 2 or ≤ 0.5, *P* value < 0.05, and variable importance in projection (VIP) ≥ 1, were considered significantly differential secondary metabolites. Significantly differentially expressed secondary metabolites were annotated using the KEGG compound database (http://www.kegg.jp/kegg/compound/), and the annotated secondary metabolites were mapped to the KEGG pathway database (http://www.kegg.jp/kegg/pathway.html).

### Insect feeding assays

The assay was conducted as described by Lewis (Lewis and van Emden, 1986). A total of 35 insects for each compound and each concentration were used for the test. The third instar *S. frugiperda* larvae of artificial rearing were fed with artificial diet (CK) (Jin et al., 2020), artificial diet supplemented with 0.02469 % phenoxyacetic acid, 0.0401 % 4-methoxybenzaldehyde, 0.0208 % apigenin and 0.0056 % 12-hydroxyoctadecanoic acid. The third instar *S. frugiperda* larvae of artificial rearing were fed with artificial diet, artificial diet with half of maize powder and artificial diet with half of maize powder supplemented with 1 % L-tryptophan. The concentrations of the test compounds were referenced to their respective proportions of all detected compounds. The body weight of the third– and sixth-instar *S. frugiperda* larvae were counted, and the growth period, pupation rate and male-to-female ratio were recorded. Compounds phenoxyacetic acid (≥ 98 %), 4-methoxybenzaldehyde (≥ 99 %), apigenin (≥ 98 %), 12-hydroxyoctadecanoic acid (≥ 85 %), and L-tryptophan (≥ 99 %) were purchased from Macklin.

### Insect behavioral preference assays

The assay was conducted as described by (Hu et al., 2020). Healthy *S. frugiperda* adult females, adult males, and third-instar larvae artificially reared under similar growth conditions were selected for the behavioral preference test. A total of 30 insects for each compound and each concentration were used for the test using “Y-shaped tubes” in a fume hood. The concentrations of the test compounds were referenced to their respective ratios. This ratio was obtained from the relative amounts of compounds in resistant line compared to the relative amounts of compounds in susceptible line. The ratio ranged from 1 to 30 and was approximately 5–10 times more than the relative amounts of compounds in susceptible line. The test concentrations were set in four groups: 1 %, 5 %, 10 % and 30 %. Prior to the test, water was placed at both ends. The insects showed no obvious reactions to the water. During testing, the two ends of the Y-tube were filled with the tested compound and water. The positions of the insects were observed and recorded every two minutes. If the insect crawled towards the compounds and remained unchanged, it was denoted as being attracted by the compound; if the insect crawled towards the water and remained unchanged, it was denoted as repelled by the compound; and if the insect showed no preference, it was denoted as no response. Compounds dodecane (≥ 99 %), octanal (≥ 99 %), methyl salicylate (≥ 99 %), 1-octen-3-ol (≥ 99 %), benzene, 1-ethenyl-4-methoxy– (≥ 98 %), *p*-cymene (≥ 98 %), hexadecanal (≥ 97 %), and methyl anthranilate (≥ 96 %) were purchased from Macklin and TCI.

### *S. frugiperda* RNA-seq data analyses

The insect samples were sent to Beijing Genomics institution (China) Co., Ltd for RNA sequencing. The high-quality RNA were used to construct a sequencing library and sequenced using an DNBSEQ-T7 (2× 150-bp read length). The raw data of transcriptome were filtered through fastp (v 0.23.1), and the filtered clean reads were separately mapped to the *S. frugiperda* coding sequence (CDS) by using Bowtie2 (v 2.2.5) for subsequent analyses (Gui et al., 2022). The obtained sam files were converted into bam files, sorted using Samtools (v1.7). We calculated the transcripts per million (TPM) value of transcripts by Salmon (v1.8.0) and identified the differentially expressed genes (DEGs) of different comparison groups with the criteria of fold change ≥ 1 and FDR ≤ 0.05 through edgeR (v 3.36.0). Kyoto Encyclopedia of Genes and Genomes (KEGG) – Automatic Annotation Server (KAAS) (https://www.genome.jp/tools/kaas/) was used to annotation DEGs. KEGG enrichment analyses were performed using and OmicShare tools (https://www.omicshare.com/tools).

### Analysis methods

TBtools (v1.100) was used to perform clustering and heat map analyses. Each compound category was normalized using TBtools (v1.100), and Origin (2022b) was used to fit each compound category to obtain trend lines. The chi-square test and independent samples *t*-test were used to analyze the significance of the insect test results. Origin (2022b) was used to generate all types of graphics.

## Acknowledgements

The authors acknowledge Dr. Xiping Yang for insightful discussions during the experiments. The authors acknowledge the sugarcane breeding team of Guangxi University for providing sugarcane materials.

## Author contributions

YZ designed the study. LS performed the experiments and analyses. LS, BC and YZ wrote the manuscript. GL-assisted sampling and plant management. CH helped manage the plants in the greenhouse and insects. MZ and CW assisted with data analysis. MF directed the microscopic observations.

## Supporting Information

**Figure S1** Susceptibility of S-115 and R-111 to *S. frugiperda* attack.

**Figure S2** Data Evaluation results of NOCs.

**Figure S3** Integral correction diagram for quantitative analysis of NOCs.

**Figure S4** Data Evaluation results of VOCs.

**Figure S5** Changing trends at different time points before and after insect infestation and the composition of compound categories for each trend.

**Table S1** Observed morphological characteristic changes in the after *S. frugiperda* infested S-115 and R-111 sugarcane lines.

**Table S2** Information of all NOCs in R-111 and S-115 (Relative amount).

**Table S3** Information of plant NOCs in R-111 and S-115 (Relative amount).

**Table S4** Difference in plant NOCs in R-111 and S-115 in healthy state (Relative amount).

**Table S5** Difference in plant NOCs between health and oral-secretion treatment in R-111 (Relative amount).

**Table S6** Difference in plant NOCs between health and oral-secretion treatment in S-115 (Relative amount).

**Table S7** Difference in plant NOCs between R-111 and S-115 by oral-secretion treatment (Relative amount).

**Table S8** Difference in plant NOCs by mechanical damaged (Relative amount).

**Table S9** Information of all VOCs in R-111 and S-115 (Relative amounts).

**Table S10** Information of plant VOCs in R-111 and S-115 (Relative amounts).

**Table S11** Difference in plant VOCs between R-111 and S-115 in healthy state (Relative amounts).

**Table S12** Difference in plant VOCs between healthy and oral-secretion-treated R-111 (Relative amounts).

**Table S13** Difference in plant VOCs between healthy and oral-secretion-treated S-115 (Relative amounts).

**Table S14** Difference in plant VOCs between R-111 and S-115 after oral-secretion treatment (Relative amounts).

**Table S15** Difference in plant VOCs after mechanical damage (Relative amounts).

## Funding

This work was supported by the National Natural Science Foundation of China [Grant No. 32100606]; Specific Research Project of Guangxi for Research Bases and Talents [GK AD21075011]; Guangxi Natural Science Foundation [GK AD20297064]; State Key Laboratory for Conservation and Utilization of Subtropical Agro-Bioresources [SKLCUSA-a202206]; Innovation Project of Guangxi Graduate Education [YCSW2022080].

## Conflicts of interest

The authors declare no competing interests.

## Data availability

The generated raw sequence data were deposited to NCBI Sequence Read Archive (SRA) database under the Bio Project Accession No PRJNA912140. The data that supports the findings of this study are available in the supporting information of this article.

